# Trophoblast survival signaling during human placentation requires HIF-induced transcription of HSP70

**DOI:** 10.1101/043851

**Authors:** Chandni V. Jain, Philip Jessmon, Charbel T. Barrak, Alan D. Bolnik, Brian A. Kilburn, Michael Hertz, D. Randall Armant

**Affiliations:** Departments of Phyiology, Wayne State University School of Medicine, Detroit, Michigan, USA; Obstetrics and Gynecology, Wayne State University School of Medicine, Detroit, Michigan, USA; Anatomy and Cell Biology, Wayne State University School of Medicine, Detroit, Michigan, USA

**Keywords:** trophoblast, hypoxia, matrix metalloproteinases, HBEGF, heat shock proteins, apoptosis

## Abstract

Survival of trophoblast cells in the low oxygen environment of human placentation requires metalloproteinase-mediated shedding of HBEGF and downstream signaling. A matrix metalloproteinase (MMP) antibody array and quantitative RT-PCR revealed upregulation of MMP2 post-transcriptionally in human first trimester HTR-8/SVneo trophoblast cells and placental explants exposed to 2% O_2_. Specific MMP inhibitors established the requirement for MMP2 in HBEGF shedding and upregulation. Hypoxia inducible factors, HIF1A and EPAS1 (HIF2A), accumulated at 2% O_2_, and HIF target genes were identified by next-generation sequencing of RNA from trophoblast cells cultured at 2% O_2_ for 0, 1, 2 and 4 hrs. Of nine genes containing HIF-response elements upregulated at 1 hour, only HSPA6 (HSP70B’) remained elevated after 4 hours. The HSP70 chaperone inhibitor VER155008 blocked upregulation of both MMP2 and HBEGF at 2% O_2_, and increased apoptosis. However, both HBEGF upregulation and apoptosis were rescued by exogenous MMP2. We propose that MMP2-mediated shedding of HBEGF, initiated by HSP70, contributes to trophoblast survival at the low O_2_ levels encountered during the first trimester, and is essential for successful pregnancy outcomes.

**Summary Statement:** Trophoblast survival during human placentation, when oxygenation is minimal, required HIF-induced transcription of HSP70, which mediated MMP2 accumulation and the transactivation of anti-apoptotic ERBB signaling by HBEGF shedding.

## INTRODUCTION

In humans, the placenta is not fully perfused with oxygenated maternal blood until after the tenth week of pregnancy, rendering the implantation site a low O_2_ environment (Burton, 2009; Burton et al., 1999; Jauniaux et al., 1991). Atypical of most cells, human trophoblast (TB) cells not only survive at low O_2_, but also proliferate more rapidly (Genbacev et al., 1996; Genbacev et al., 1997; Kilburn et al., 2000). The epidermal growth factor (EGF) family member, heparin-binding EGF-like growth factor (HBEGF) is highly expressed in the uteroplacental compartment during implantation and early placentation (Jessmon et al., 2009). We previously reported that HBEGF prevents death of human trophoblast cells exposed to low O_2_ concentrations (Armant et al., 2006). Although HBEGF is upregulated ~100-fold in TB cells after exposure to 2% O_2_, HBEGF mRNA remains unchanged and is abundant (2500 copies/cell) at both O_2_ concentrations (Armant et al., 2006). Survival of TB cells at low (2%) O_2_ requires metalloproteinase-mediated shedding of membrane-bound proHBEGF and HBEGF downstream signaling (Armant et al., 2006). HBEGF is synthesized as transmembrane proHBEGF, and secreted through shedding from the plasma membrane by proteolytic cleavage, allowing it to activate ERBB receptor tyrosine kinases, including its cognate receptors, EGF receptor/ERBB1 and ERBB4 (Holbro and Hynes, 2004). As O_2_ increases with the full perfusion of the intravillous space after 10 weeks of gestation (Burton, 2009; Burton et al., 1999; Jauniaux et al., 1991), HBEGF positively regulates TB motility and invasion (Jessmon et al., 2010; Leach et al., 2004). The hypertensive pregnancy disorder, preeclampsia, in which TB invasion is reduced and apoptosis elevated, is characterized by a reduction in HBEGF and other components of the EGF signaling system (Armant et al., 2015; Jessmon et al., 2009; Leach et al., 2002). It can, therefore, be hypothesized that HBEGF production could be a factor in trophoblast dysfunction associated with placental insufficiency disorders.

In human TB cells, HBEGF activates p38 (MAPK14), ERK (MAPK1), and JNK (MAPK8) (Jessmon et al., 2010). During hypoxia, HBEGF upregulates its biosynthesis through an autocrine feedback mechanism, and prevents apoptosis through a parallel signaling pathway (Armant et al., 2006; Jessmon et al., 2010). Activation of ERBB receptor tyrosine kinase by HBEGF prevents apoptosis through a mechanism mediated by p38 MAPK, while autocrine upregulation of HBEGF appears to be mediated by any one of the MAPKs, p38, ERK, or JNK, based on experiments using specific inhibitors (Jessmon et al., 2010). However, it is unclear whether the MAPKs specifically function downstream of HBEGF in its autocrine upregulation, or also operate further upstream in the cascade initiated by hypoxia to regulate proHBEGF shedding.

It is well established that metalloproteinases contribute to invasion and tissue remodeling (Loffek et al., 2011; Stamenkovic, 2003). Successful implantation and TB invasion are closely linked to the expression of matrix metalloproteinases (MMPs) that degrade basement membranes (Librach et al., 1991). The gelatinases (gelatinase A/MMP2; 72-kDa, and gelatinase B/MMP9; 92-kDa), which target major components of basement membranes (e.g., collagen IV), are expressed by TB cells, and therefore, regarded as key to the invasion process (Isaka et al., 2003). MMP2 and MMP9 are differentially expressed in first trimester TB cells, with MMP2 more prominently secreted until 9 weeks (Staun-Ram et al., 2004; Xu et al., 2000). MMP2 is constitutively expressed in TB cells throughout pregnancy, but its activity is diminished in the full-term placenta. MMP9 is mainly expressed by TB cells during the period after Week 9, and also decreases at term. Thus, MMP2 and MMP9 are expressed and functioning throughout the periods of low and then elevated O_2_ in the reproductive tract, characterized by high TB proliferation and then elevated invasion, respectively (Jovanovic et al., 2010; Librach et al., 1991). While their role in tissue remodeling is well known (Loffek et al., 2011; Stamenkovic, 2003), emerging evidence reveals that MMPs also participate in shedding of membrane-anchored signaling molecules, which contributes to chemokine and growth factor activation (Burton, 2009; Chow and Fernandez-Patron, 2007; Powell et al., 1999).

Shedding is mediated by membrane-type MMPs and MMPs bound to membrane receptors, including MMP2 and MMP9 (Sternlicht and Werb, 2001). Cheng et. al reported that bradykinin-induced proliferation of rabbit corneal cells is blocked by an inhibitor of both MMP2 and MMP9, as well as an inhibitor of HBEGF, suggesting that an MMP could contribute to the cleavage of proHBEGF (Cheng et al., 2012). MMP2 participates in a proteolytic cascade in αT3-1 cells that directs both proHBEGF shedding and EGFR transactivation (Roelle et al., 2003). In addition, its rapid release in CMTC9 cells after E2 stimulation is correlated with cleavage of proHBEGF (Torres et al., 2009). We previously found that treatment with a general metalloproteinase inhibitor blocks HBEGF accumulation in human TB cells at low O_2_, causing apoptosis that is rescued with recombinant HBEGF, but not other EGF-like ligands (Armant et al., 2006). Autocrine HBEGF activity in TB cells requires, in addition to metalloproteinase-mediated shedding of HBEGF, binding to either EGFR or ERBB4. Blocking HBEGF signaling prevents its upregulation at 2% O_2_, suggesting that low levels of resident proHBEGF are cleaved through activation of metalloproteinases to initiate its autocrine accumulation downstream of ERBB receptors in a positive feedback loop.

To understand the molecular mechanism of TB survival at low O_2_, the HTR-8/SVneo human TB cell line (Graham et al., 1993) was used. The HTR-8/SVneo cell line is an immortalized cell line (Graham et al., 1998) that was established from human first-trimester cytotrophoblast cells (Graham et al., 1993). HTR-8/SVneo cells express cytokeratin (KRT7), a marker for trophoblast and other epithelial cells, as well as the beta subunit of human chorionic gonadotropin (β-hCG), which is trophoblast-specific (Kilburn et al., 2000). This cell line possess the ability to invade Matrigel basement membrane without being tumorigenic, and expresses the trophoblast-specific major histocompatibility protein, HLA-G, when induced by Matrigel differentiate (Kilburn et al., 2000). In response to Matrigel, HTR-8/SVneo cells switch integrin expression (Damsky) in association with invasive differentiation, while in response to hypoxia, proliferation increases and extravillous differentiation is inhibited (Kilburn et al., 2000). To examine the hypothesis that MMPs contribute to the shedding and upregulation of HBEGF at low O_2_ concentrations required for TB survival in the first trimester, we used HTR-8/SVneo human TB cells, with confirmatory experiments using first trimester villous explants (Castellucci et al., 1990; Genbacev et al., 1992).

## RESULTS

### Upregulation of MMP2 at low O_2_

Using a human MMP antibody array, we observed that relative expression of MMP2, but not MMP9 or other MMPs, increased 1.7-fold in HTR-8/SVneo TB cells cultured for 4 hrs at 2% O_2_ (Fig. 1A), as compared to TB cells cultured at 20% O_2_. The specific increase in expression of MMP2, and not MMP9, was confirmed by western blotting (Fig. 1B). Cellular MMP2 was quantified temporally by ELISA in TB cells after shifting to 2% O_2_. The MMP2 concentration abruptly increased 164-fold (p<0.0001) after 2 hours of culture at 2% O_2_ (Fig. 1C). These findings indicate that low O_2_ upregulates MMP2 prior to the observed increase in HBEGF, which occurs at 4 hrs (Armant et al., 2006).

**Fig. 1.**
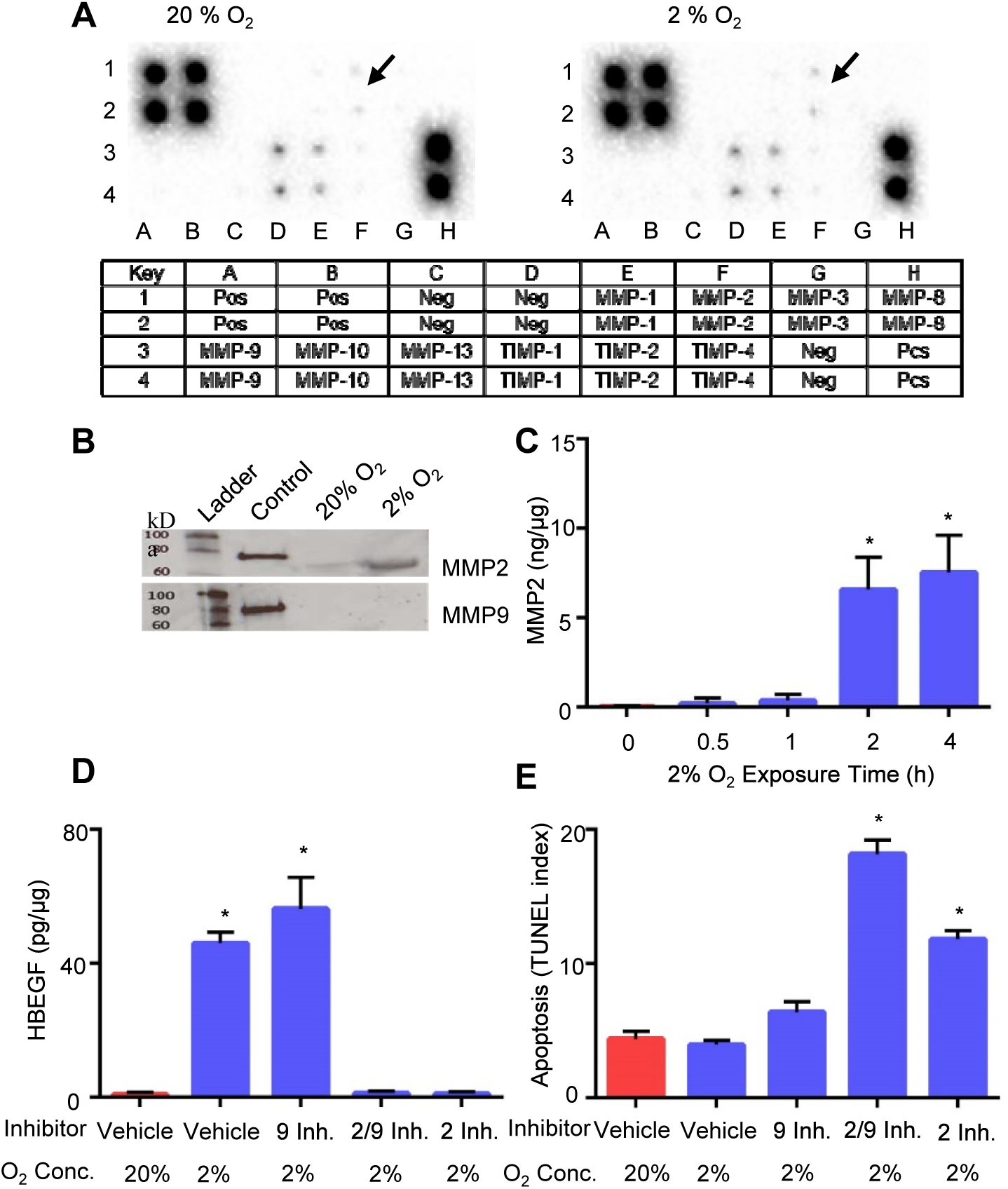
Upregulation of MMP2 at low O_2_, and its effects on HBEGF and cell survival. A. Expression arrays for MMP-related proteins incubated with extracts of HTR-8/SVneo cells cultured at 20% (left) or 2% (right) O_2_. Arrows indicate elevated MMP2 at 2% compared to 20% O_2_. The key below indicates the location of duplicate antibody probes for each protein according to the coordinates shown, including both positive (Pos) and Negative (Neg) controls. B. Western blots of molecular weight standards (Ladder), recombinant MMP2 or MMP9 (Control), and extracts of TB cells cultured at 20% or 2% O_2_, as indicated. The upper blot was labeled with anti-MMP2, and the lower blot was labeled with anti-MMP9. C. MMP2 quantified by ELISA in TB cells cultured 0-4 h at 2% O_2_. D. HBEGF quantified by ELISA in TB cells treated during culture for 4 h with the indicated MMP inhibitor or vehicle, and concentration of O_2_. E. Apoptosis was quantified in TB cells cultured as in D using the TUNEL assay. *, p<0.05, compared to the control (0 h, vehicle/20% O_2_).

### Role of MMP2 in HBEGF upregulation

To determine whether MMP2 has a functional role in HBEGF regulation by O_2_, specific inhibitors of MMP2 and MMP9 were supplemented in culture medium during manipulation of O_2_ levels. HBEGF failed to accumulate in TB cells cultured at 2% O_2_ for 4 h when MMP2 was inhibited (Fig. 1D). However, MMP9 inhibition had no effect on the normal accumulation of HBEGF at low O_2_. A third inhibitor that targets both MMP2 and MMP9 also inhibited HBEGF accumulation. Additionally, TUNEL assays showed increased (p<0.0001) TB cell death when both MMP2 and MMP9, or only MMP2, were inhibited (Fig. 1E). Inhibition of only MMP9 produced apoptosis levels similar to control. We conclude that MMP2, but not MMP9, either directly or indirectly, mediates the proteolytic cleavage of proHBEGF, and subsequent survival signaling.

### Role of MAPKs in MMP2 upregulation

We previously showed that upregulation of HBEGF at low O_2_ requires MAPK signaling (Jessmon et al., 2010). MMP2 was quantified by ELISA in extracts of TB cells cultured at 2% O_2_ for 4 hours, with or without specific inhibitors of ERK, p38 and JNK, or the corresponding inactive structural analogs (Table 1). Inhibitors of MAPKs had no effect on the increased MMP2 expression at 2% O_2_, suggesting that the upregulation of MMP2 is independent of MAPK, and that MAPK signaling most likely functions exclusively downstream of ERBB activation.

**Table 1.** MMP2 upregulation is independent of MAPKs

| **Table 1. MMP2 upregulation is independent of MAPKs** |  |  |
| --- | --- | --- |
| **% O<sub>2</sub>** | **Treatment** | **MMP2 (ng/ml)** |
| 20 | Vehicle | 0.03 ± 0.03 |
| 2 | Vehicle | 7.27 ± 0.74 * |
| 2 | U 0126/ERK inhibitor | 6.07 ± 0.55 * |
| 2 | U0124/ERK Negative control | 7.40 ± 0.66 * |
| 2 | JNK inhibitor | 5.44 ± 1.98 * |
| 2 | JNK Negative control | 6.53 ± 1.23 * |
| 2 | SB203580 P38 inhibitor | 6.17 ± 0.87 * |
| 2 | SB 202474 P38 Negative control | 7.15 ± 0.89 * |
| 2 | All 3 inhibitors | 5.86 ± 1.51 * |
| 2 | All 3 neg. controls | 6.34 ± 0.68 * |

\* p<0.0001

### MMP2 mRNA expression

To determine whether MMP2 is transcriptionally upregulated at low O_2_, its expression was quantified by qPCR, using RNA extracted from HTR-8/SVneo cells cultured at 20% and 2% O_2_ for 1-4 hrs. Expression values, normalized to GAPDH, were calculated for both MMP2 (Fig.2A) and HBEGF (Fig. 2B). MMP2 mRNA, like HBEGF message, did not change (p=0.8) at 2% O_2_, suggesting that their proteins are both regulated post-transcriptionally.

### Role of transcription in HBEGF upregulation

TB cells were cultured at 2% O_2_ with or without α-amanitin, an inhibitor of RNA polymerase-II (Lindell et al., 1970; Stirpe and Fiume, 1967), to examine the role of *de novo* transcription in the upregulation of HBEGF. Elevation of HBEGF protein after culture at 2% O_2_ for 6 h was abrogated in a dose-dependent manner by α-amanitin, with maximal inhibition attained at 5 μg/ml (Fig. 2C). CoCl_2_, a chemical mimic of hypoxia that stabilizes hypoxia inducible factor (HIF) (Jiang et al., 1997), was used to examine its effect on HBEGF. At 20% O_2_, CoCl_2_ increased (p<0.0001) HBEGF levels. As expected, CoCl_2_ induced accumulation of HIF1A and HIF2A, as observed by western blotting (Fig. 2E), similar to the effect of low O_2_. α-Amanitin also blocked the CoCl_2_-mediated increase in HBEGF (Fig. 2D). The effects of α-amanitin on HBEGF suggest that transcription of genes, perhaps those regulated by HIF1A or HIF2A, is required to initiate HBEGF shedding and accumulation.

**Fig. 2.**
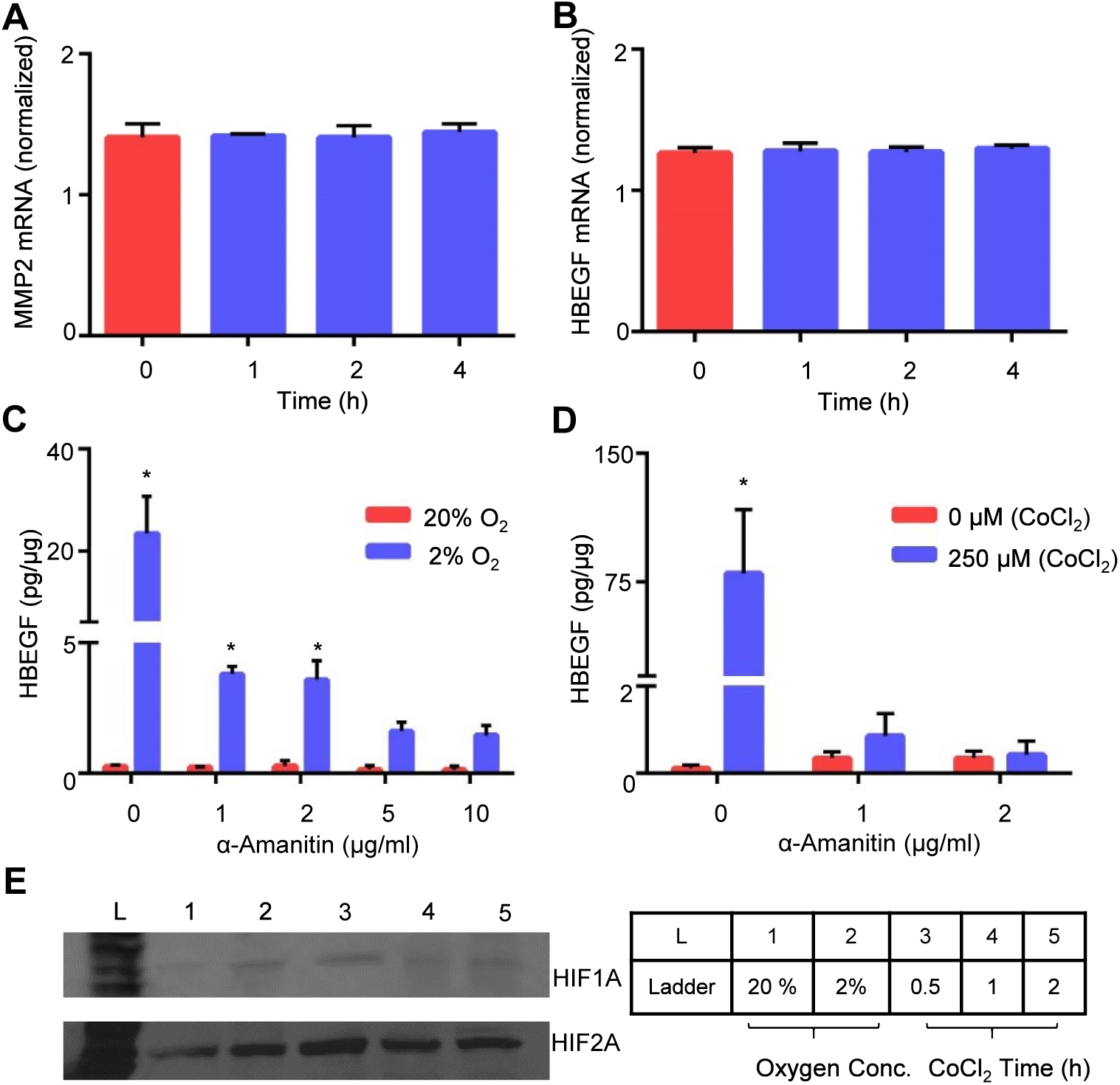
Expression of MMP2 and HBEGF at 2% O_2_ in HTR-8/SVneo cells. MMP2 (A) and HBEGF (B) mRNA were measured by qPCR after culture for indicated time at 2% O_2_. C. HBEGF protein measured by ELISA in extracts of TB cells cultured at the indicated concentrations of α-amanitin and O_2_. D. HBEGF protein measured by ELISA in extracts of TB cells cultured at the indicated concentrations of α-amanitin and CoCl_2_ at 20% O_2_. * p < 0.05 compared to no treatment control. E. Western blots of HIF1A (~ 93 kDa) and HIF2A (~96 kDa) in lysates of TB cells treated at 20% O_2_ or 2% O_2_ for 4h, or at 20% O_2_ in the presence of 250 μM CoCl_2_ for 0.5 - 2 h, as indicated in the Key to the right.

### Identification of differentially expressed transcripts

Transcripts differentially expressed in TB cells at the two O_2_ levels were identified, using a non-biased NGS approach. RNA-seq analysis using the GGA revealed a total of 9 upregulated (Table 2) and 120 downregulated genes (**Table S1**) after 1 hr of exposure to 2% O_2_ (Fig. 3A). Of the 9 upregulated genes, HSPA6 was the most dramatically upregulated (870-fold) (Table 2), and the only gene that remained elevated at 2 h (Fig. 3B) and 4 h (Fig. 3C). The increase in HSPA6 expression at 2% O_2_ was validated using qPCR and western blotting (Fig. 3D,E). Extraction of the promoter region using the GGA MatInspector (Cartharius et al., 2005) program revealed two overlapping HRE’s 2.4 kbp upstream of the 5’-most HSPA6 transcriptional start site (**Fig. S1**), suggesting that HSPA6 could operate downstream of hypoxia to regulate MMP2 and HBEGF biosynthesis at low O_2_.

**Table 2.**
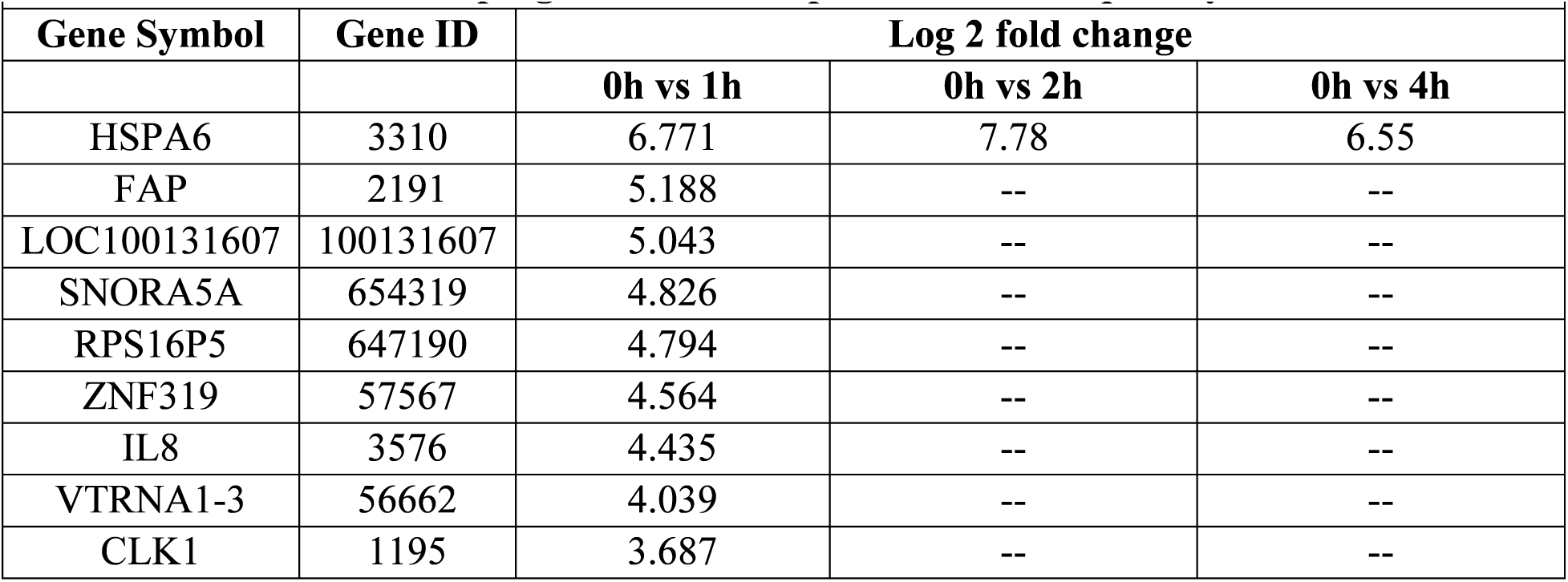
Upregulated transcripts from RNASeq Analysis

| Table 2. Upregulated transcripts from RNASeq Analysis |  |  |  |  |
| --- | --- | --- | --- | --- |
| Gene Symbol | Gene ID | Log 2 fold change |  |  |
|  |  | 0h vs 1h | 0h vs 2h | 0h vs 4h |
| HSPA6 | 3310 | 6.771 | 7.78 | 6.55 |
| FAP | 2191 | 5.188 | -- | -- |
| LOC100131607 | 100131607 | 5.043 | -- | -- |
| SNORA5A | 654319 | 4.826 | -- | -- |
| RPS16P5 | 647190 | 4.794 | -- | -- |
| ZNF319 | 57567 | 4.564 | -- | -- |
| IL8 | 3576 | 4.435 | -- | -- |
| VTRNA1-3 | 56662 | 4.039 | -- | -- |
| CLK1 | 1195 | 3.687 | -- | -- |

**Fig. 3.**
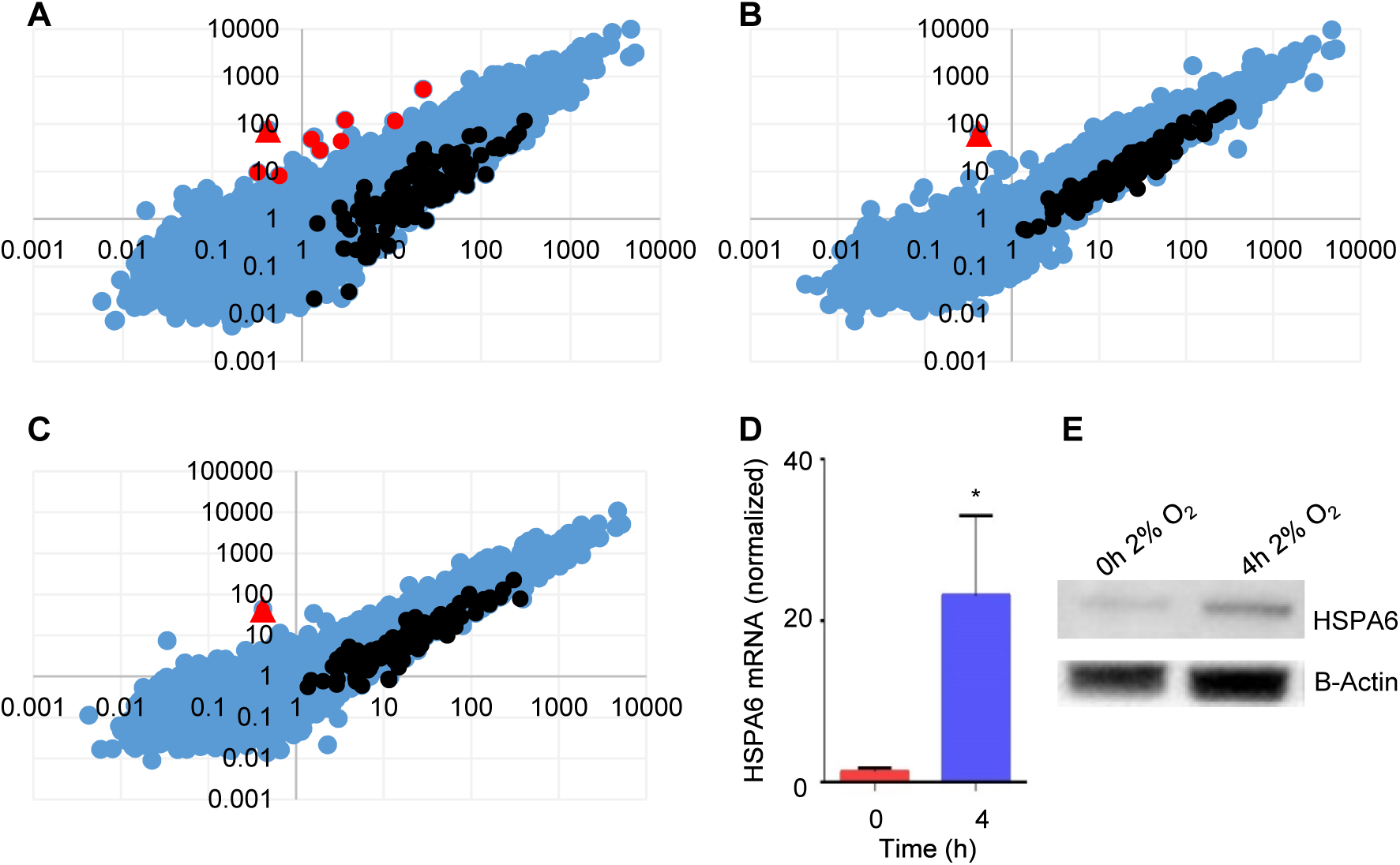
Transcriptome analysis of TB cells exposed to 2% O_2_. A-C. Abundance of transcripts compared using GGA work station between cells cultured at 2% O_2_ for 0 h (x-axis) and 1 h (A, y-axis), 2 h (B, y-axis) or 4 h (C, y-axis). Unchanged transcripts are indicated in blue, upregulated transcripts are red, and downregulated transcripts are black. HSPA6 (red triangle) was the most highly upregulated mRNA, and only transcript upregulated at 2 h and 4 h. D. HSPA6 mRNA was measured by qPCR in TB cells cultured for indicated time at 2% O_2_. E. Western blots of HSPA6 (~70 kDa) and β-actin (~43 kDa) in lysates of TB cells cultured for indicated time at 2% O_2_.

### HSP70 function in MMP2 and HBEGF upregulation

Inhibition of HSP70 with a pharmacological inhibitor, VER 155008 (Schlecht et al., 2013), in TB cells cultured at 2% O_2_ caused a dose-dependent decrease in MMP2 (Fig. 4A) and HBEGF (Fig. 4B). TUNEL assays showed increased (p<0.0001) TB cell death with HSP70 inhibition (Fig. 4C), suggesting that survival at 2% O_2_, through upregulation of MMP2 and HBEGF, requires HSPA6 activity. Using ELISA, we demonstrated that MMP2 was not only required for HBEGF upregulation (Fig. 1D), but that addition of rMMP2 was sufficient to increase (p<0.0001) HBEGF at 20% O_2_ to the same extent as TB culture at 2% O_2_ (Fig. 4D). Furthermore, addition of rMMP2 rescued (p<0.0001) HBEGF biosynthesis in the presence of HSP70 inhibitor at 2% O_2_ (Fig. 4D).

**Fig. 4.**
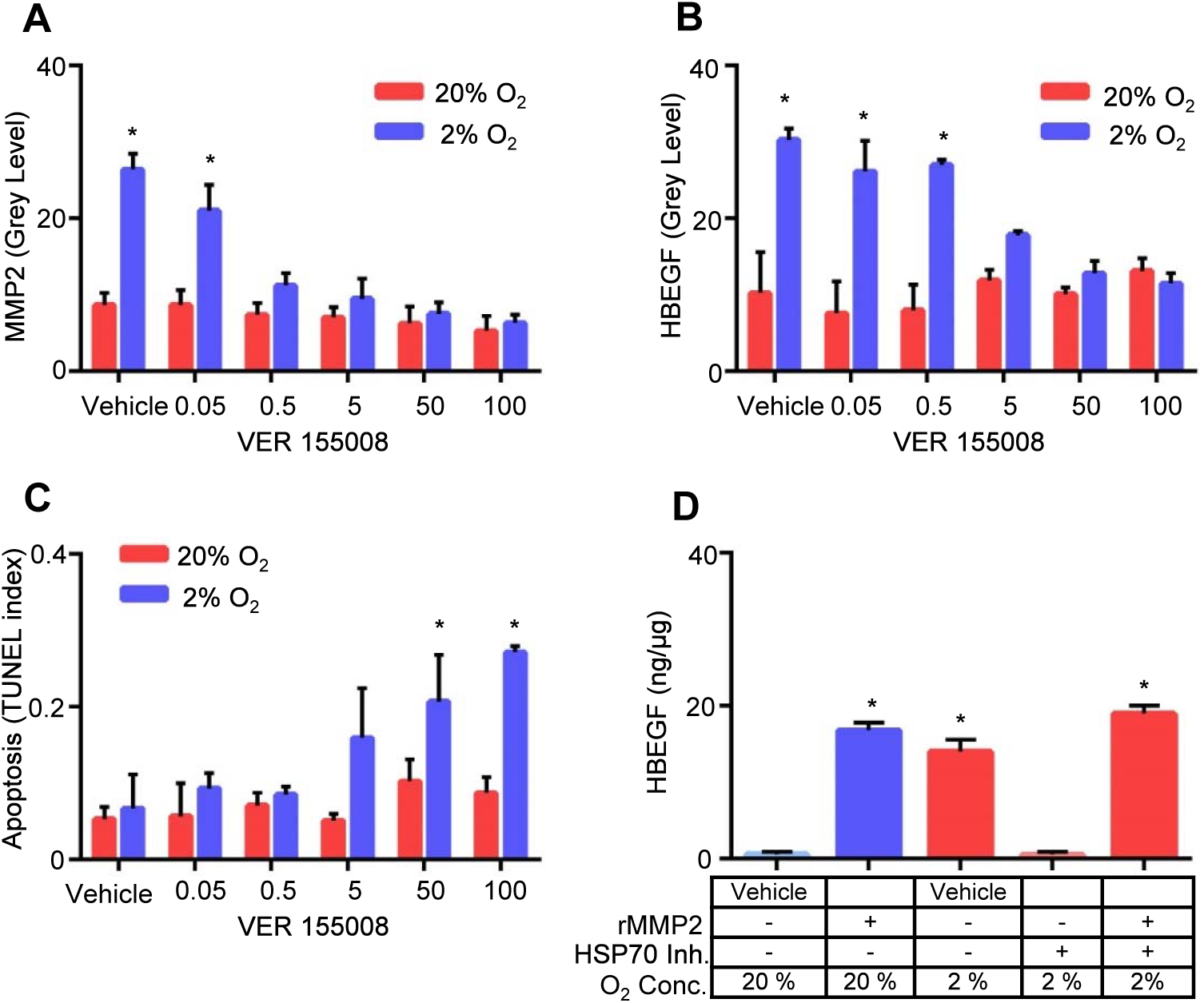
Regulation of MMP2, HBEGF and cell survival by HSP70. HSP70 was inhibited using the pharmacological inhibitor VER 155008 during culture of HTR-8/SVneo cells for 4 h at 20% or 2% O_2_. At 2% O_2_, HSP70 inhibition caused a dose-dependent decrease in both MMP2 (A) and HBEGF (B), measured by immunohistochemistry, and concomitantly increased cell death (C), measured by TUNEL. Cells cultured at 20% O_2_ were unaffected. D. ELISA for HBEGF in cellular extracts of TB cells cultured for 4 h at either 20% or 2% O_2_ in the presence of recombinant MMP2 (rMMP2) and HSP70 inhibitor (VER 155008), as indicated. *, p<0.05, compared to vehicle/20% O_2_.

These results demonstrate that upregulation of MMP2 requires HSPA6 (HSP70B’) activity, while upregulation of HBEGF requires only shedding mediated by MMP2.

### Regulation of MMP2 in first trimester explants

Using immunocytochemistry and ELISA, it was confirmed that HSP70 inhibitor prevents the upregulation of MMP2 (Fig. 5A,B) and HBEGF (Fig. 5C,D) in first trimester explants cultured at 2% O_2_. In addition, rMMP2 alone increased (p<0.0001) HBEGF at 20% O_2_, and rescued (p<0.0001) HBEGF upregulation at 2% O_2_ during treatment with HSP70 inhibitor (Fig. 5C,D). Cell death assays showed increased (p<0.05) cell death during HSP70 inhibition that was rescued (p<0.05) by rMMP2 (Fig. 5E,F), suggesting that survival at 2% O_2_, through upregulation of MMP2 and HBEGF, requires HSPA6 activity.

**Fig. 5.**
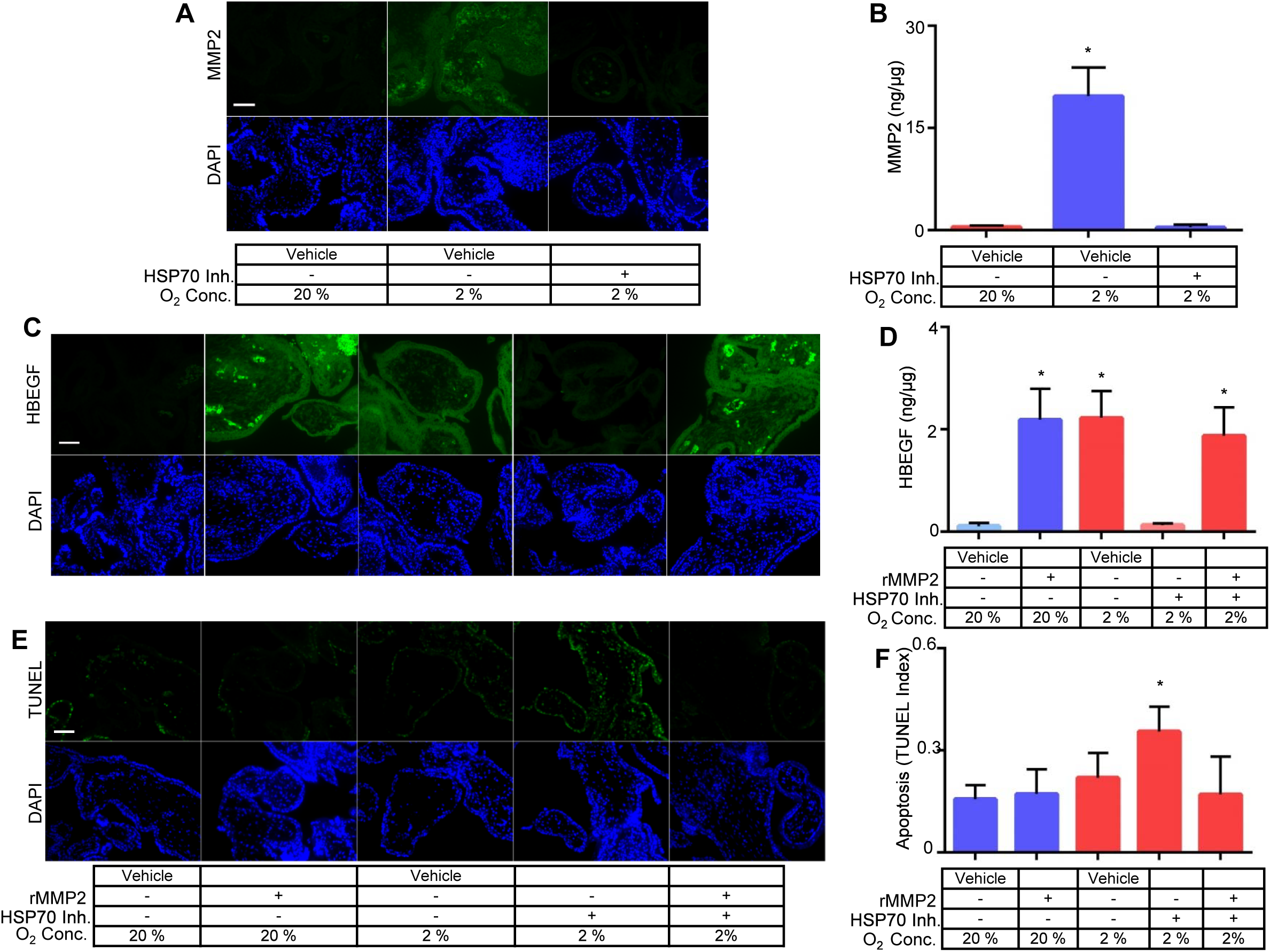
Regulation of MMP2 in first trimester chorionic villous explants. First trimester villous explants cultured for 6 h at either 20% or 2% O_2_ in the presence of recombinant MMP2 (rMMP2) and VER 155008 (HSP70 inh.), as indicated, were dually stained for DAPI and MMP2 (A), HBEGF (C), or TUNEL (E), using immunofluorescence microscopy. Size bars indicating 100 μm are shown in A, C and E. Cell extracts were assayed by ELISA for MMP2 (B) and HBEGF (D), as well as for cell death by TUNEL (F). *, p<0.0001 in B and D, p<0.05 in F, compared to vehicle/20% O_2_.

## DISCUSSION

The regulation of HBEGF signaling during implantation and placentation is critical for TB survival and invasion (Jessmon et al., 2009; Lim and Dey, 2009). An autocrine, post-transcriptional mechanism induced at low O_2_ upregulates HBEGF synthesis and secretion, providing an important survival factor during early gestation (Armant et al., 2006; Jessmon et al., 2009). The present study provides evidence that a transcriptional event is required to initiate HBEGF shedding downstream of HIF signaling. In seeking a metalloproteinase responsible for HBEGF shedding, a human MMP antibody array, verified by western blot and ELISA, implicated a protease cascade that includes MMP2. A functional role for MMP2 in the accumulation of HBEGF was established using a specific inhibitor. However, MMP2 mRNA concentration was unaltered by reduced O_2_, suggesting that it was regulated post-transcriptionally. Transcriptomic analysis identified HSPA6, a member of the HSP70 family, as a potential regulator upstream of MMP2. Using an inhibitor of HSP70 and exogenous rMMP2, evidence was obtained supporting a mechanism wherein low O_2_ concentrations, as experienced by TB cells during the first 10 weeks of placentation, induce HIF-mediated transcription of HSP70, a chaperone that directly or indirectly facilitates MMP2 accumulation. MMP2, in turn, appears to participate in a protease cascade that sheds HBEGF from the cell surface (Hitchon et al., 2002; Mohammad et al., 2010). The production of sHBEGF initiates autocrine signaling through its cognate receptors, EGFR and ERBB4, which activates its biosynthesis from constitutively-expressed, but latent, HBEGF mRNA (Fig.6).

**Fig. 6.**
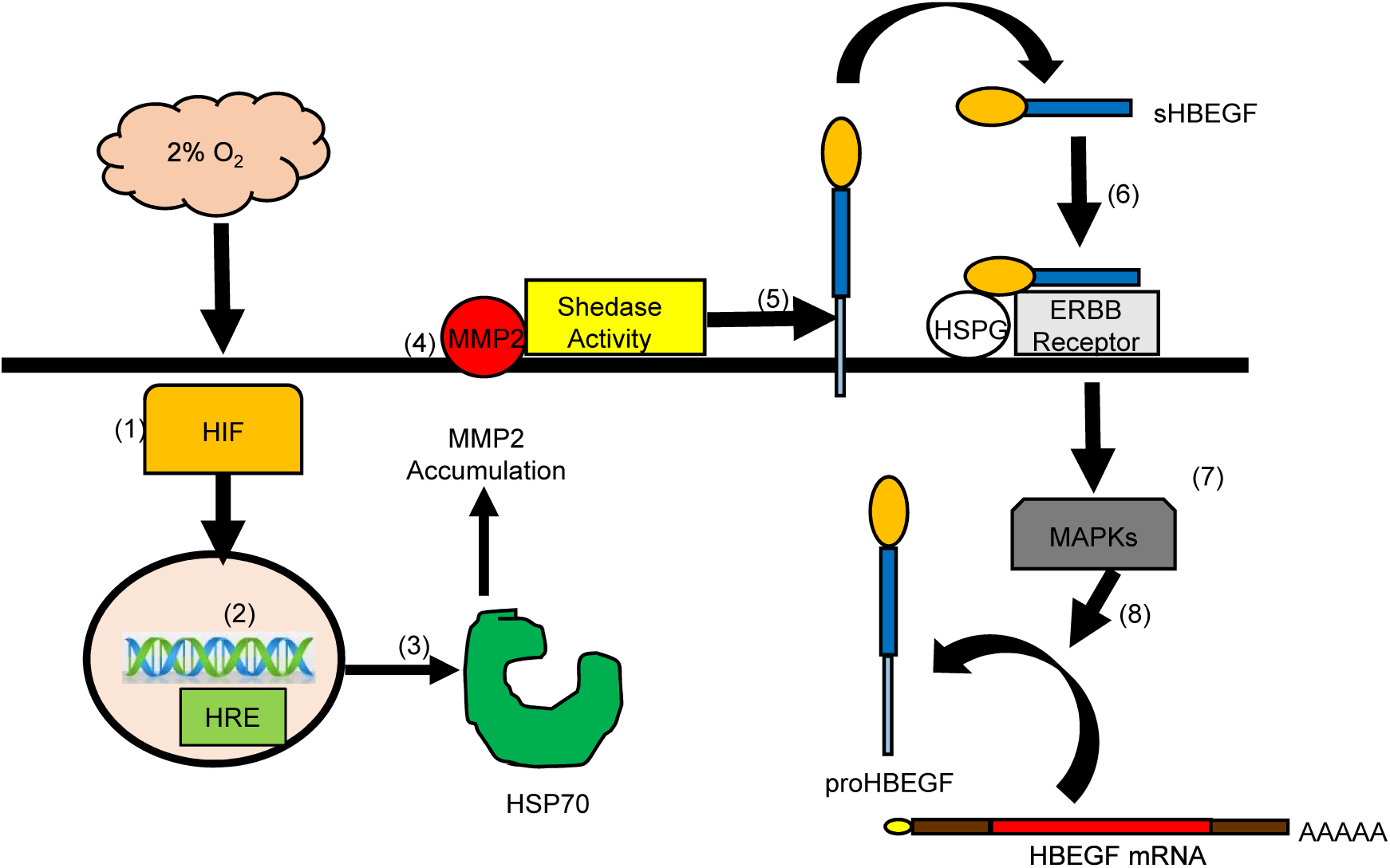
Proposed Mechanism. Low (2%) O_2_ stabilizes cytoplasmic HIF1A and HIF2A (1) that accumulate and translocate to the nucleus (2) where they dimerize to HIF1B and bind to hypoxia response elements (HRE) in the regulatory regions of target genes, including HSPA6/HSP70B’ (3). HSP70 regulates the accumulation of activated MMP2 at the cell surface (4), which is required for shedding (5) of the extracellular domain of proHBEGF. The released sHBEGF binds (6) to its receptors, ERBB1 and ERBB4, through its EGF-like domain and to heparan sulfate proteoglycans (HSPG) through its heparin-binding domain. ERBB downstream signaling activates MAPKs (7) required for biosynthesis of proHBEGF from the pool of HBEGF mRNA (8). HBEGF autocrine signaling thereby increases HBEGF secretion to achieve extracellular HBEGF concentrations (>1 nM) sufficient to inhibit apoptosis at 2% O_2_.

Supplementation with HBEGF and other EGF family members such as EGF or TGFα has little effect on proliferation rates, but is highly effective at converting TB cells to an invasive phenotype (Leach et al., 2004). The extensive expression of HBEGF in TB cells, particularly within extravillous populations (Leach et al., 2002), could be vital for their invasive activities during the establishment of pregnancy and physiological conversion of spiral arteries. Therefore, it has been established that HBEGF regulates trophoblast survival and invasion, two crucial functions that are compromised in pregnancies complicated by preeclampsia (Brosens et al., 1972; DiFederico et al., 1999). HBEGF and other components of the EGF signaling system are disrupted in placentas of women with preeclampsia (Armant et al., 2015; Leach et al., 2002). Trophoblast invasion is shallow in preeclampsia, possibly due to a lack of HBEGF-induced cell migration and a rise in apoptosis, exacerbated by reduced cytoprotection. The increased oxidative damage to trophoblast cells (Redman and Sargent, 2005) and the decrease in HBEGF in preeclamptic placentas is consistent with the hypothesis that HBEGF is an important survival factor throughout gestation.

Jessmon et. al showed that HBEGF activates ERK, p38 and JNK in human TB cells, and treatment with specific inhibitors indicated that hypoxia upregulates HBEGF biosynthesis through any one of the three examined MAPKs (Armant et al., 2006; Jessmon et al., 2010). However, it was unclear whether this MAPK pathway was functional upstream or downstream of HBEGF shedding. Therefore, MMP2 was quantified in human TB cells cultured at 2% O_2_ with specific inhibitors of ERK, p38 and JNK. These inhibitors did not influence the upregulation of MMP2 at low O_2_, suggesting that the MAPKs function only downstream of HBEGF signaling through the ERBB1/4 tyrosine kinases in human TB cells, as indicated in Fig.6.

MMP2 was not among the genes that were transcriptionally regulated at low O_2_, but nevertheless appears to participate in a proteolytic cascade that culminates in HBEGF shedding. Studies of early placentation suggest that MMP2 is important in TB invasion (Staun-Ram et al., 2004). Interestingly, very low O_2_ (0.1%) increases MMP2 mRNA levels 1.7-fold in TB cells isolated from first trimester placentae (Onogi et al., 2011), which was not found in our study. However, culture at 2% O_2_ could differ significantly from 0.1% O_2_, and account for the different outcomes. Using specific inhibitors and rMMP2, we showed that MMP2, but not HBEGF, is regulated by HSP70. It appears that HSP70 stabilizes either MMP2 or a protein regulating its accumulation at the cell surface for shedding of HBEGF.

Although both MMP2 and HBEGF appear to be regulated by O_2_ post-transcriptionally, HBEGF upregulation by low O_2_ or CoCl_2_ was blocked by α-amanitin, suggesting the requirement for transcription. HIF proteins are stabilized at low O_2_ (Wang and Semenza, 1993), and activate transcription of targeted genes by interaction with specific HREs in the gene promoters (Semenza, 2003), along with transcriptional coactivators to form an initiation complex (Lisy and Peet, 2008). We suspected that either HIF1A or HIF2A could be involved in the response of TB cells to low O_2_, based on their accumulation under the influence of CoCl_2_ or 2% O_2_, which was previously reported in JEG3 choriocarcinoma cells (Park et al., 2014). Interestingly, the transcriptome analysis revealed HSPA6/HSP70B’ as the most highly upregulated gene at 1-4 h, while no changes occurred in transcripts of metalloproteinases. Extraction of the HSPA6 promoter revealed two overlapping HREs on the sense and anti-sense strands. HREs also appeared in the promoters of the eight other mRNAs that increased 1 h after exposure of TB cells to low O_2_. In rabbit chondrocytes and JEG3 cells increased expression of HIF1A under hypoxia (2%) or simulated hypoxia (CoCl_2_) resulted in an increase in HSP70 expression that was ablated by HIF1A inhibition or knockdown (Park et al., 2014; Tsuchida et al., 2014). These data suggest that HSP70 transcription is induced by HIF1A signaling.

Induction of heat shock proteins (HSPs) in response to cellular stress has been proposed as a potential strategy of eukaryotic cells to combat lethal conditions (Ananthan et al., 1986; Goff and Goldberg, 1985; Parsell and Sauer, 1989). Within the HSP family, HSP70 functions in the recovery of cells from stress, and in guarding against further insults (Fulda et al., 2010). HSP70 is a multigene family that includes the stress-inducible members HSPA6 and HSPA1A/HSP70-1 (Schwarz et al., 2003; Tavaria et al., 1996). HSP70 protects stressed cells by recognizing nascent polypeptides, unstructured regions of protein, and exposed hydrophobic stretches of amino acid (Nollen and Morimoto, 2002). The binding of exposed hydrophobic residues in unfolded or partially folded proteins is regulated by ATP-hydrolysis-induced conformational changes in the ATPase domain of HSP70, which is stimulated by co-chaperones (Nollen and Morimoto, 2002). HSP70 is transcribed in response to stress due to activation of heat shock transcription factor (HSF) (Abravaya et al., 1992). Abravaya et.al proposed a model suggesting that physiological stress creates missfolded proteins that compete with HSF for binding to HSP70, resulting in release of transcriptionally active, DNA-binding HSF to induce transcription of heat shock genes. Baird et.al has shown that this HSF-mediated transcription is increased under low O_2_ due to direct binding of HIF1A to its HRE on the HSF intron (Baird et al., 2006). These reports are consistent with a mechanism wherein HSP70 transcription is mediated by HSF or HIF1A at low O_2_ in TB cells.

HSP70 forms a molecular chaperone complex with HSP90, another highly conserved and essential HSP family member (Pratt et al., 2015; Pratt and Toft, 2003). This molecular chaperone complex is dependent on the hydrolysis of ATP, and ADP/ATP exchange, and is mediated by association with HOP (Hsp70/Hsp90-organising protein)/STIP1 (stress-induced phosphoprotein 1) (Carrigan et al., 2006; Chen and Smith, 1998). The HSP70/HSP90 molecular chaperone complex, optimized by HOP, coordinates interactions that ensure folding and conformational regulation of proteins under stress (Hernandez et al., 2002; Wegele et al., 2004). RNAi knockdown of HOP in pancreatic cancer cells decreases expression of MMP2 (Walsh et al., 2011). Additionally, inhibition or depletion of HSP70 in the breast cancer cell line, MA-MB-231, decreases cell migration and invasion due to a lack of MMP2 activation (Sims et al., 2011). This suggests that molecular relationships between HSPs, various signaling proteins, and partner proteins mediate the integrity of signal transduction pathways (Nollen and Morimoto, 2002). Our data further supports the role for HSP70 in mediating MMP2 function, which is required in TB cells for HBEGF survival signaling.

The signaling cascade delineated in the present study (Fig. 6) could influence multiple physiological processes that play a key role in placentation. There is substantial evidence to support that HBEGF expression is altered in preeclampsia, and that its protein levels are decreased in preeclamptic placentas (Armant et al., 2015; Leach et al., 2002). MMP2 belongs to a family of extracellular matrix-remodeling enzymes that have been implicated in the regulation of vasculogenesis, which is disrupted in placental insufficiency disorders, such as preeclampsia. There is compelling evidence that MMP2, but not MMP9, increases in women who subsequently develop preeclampsia (Eleuterio et al., 2015; Montagnana et al., 2009; Myers et al., 2005; Narumiya et al., 2001). HSP70 also appears to be altered in adverse pregnancies. In a pilot study, higher levels of HSP70 were reported in patients with early onset of severe PE (Jirecek et al., 2002; Yung et al., 2014). Fukushima et.al reported that serum levels of HSP70 were constant throughout normal pregnancy, but increased significantly in women with preeclampsia (Fukushima et al., 2005; Molvarec et al., 2006) or preterm delivery (Fukushima et al., 2005). Increased circulating HSP70 in preeclamptic patients could be due to systemic inflammation as a result of disease and increased oxidative stress (Ekambaram, 2011; Molvarec et al., 2009). In term preeclamptic placentas, HIF1A and HSP70 are both elevated and, localized prominently in syncytiotrophoblasts and villous endothelial cells (Park et al., 2014). In another study of HSP70 in term placentas, both mRNA and protein increased in women with preeclampsia and intrauterine growth restriction (Liu et al., 2008). However, there has been no information reported on the expression or role of placental HSP70 in the first trimester prior to this study.

Using an established TB cell line and a villous explant model, we have established a role for HSPA6 (HSP70B’), a HIF-regulated gene, in the regulation of MMP2 required for HBEGF shedding at low O_2_. These findings suggest that TB survival in the low O_2_ environment during early pregnancy requires this signaling pathway. Disruption of any component during the first trimester could compromise TB survival and function, leading to placental insufficiency and the resulting obstetrical complications of pregnancy.

## MATERIALS AND METHODS

### Cell Culture and Treatments

The first trimester human TB cell line, HTR-8/SVneo (Graham et al., 1993), were grown in either 96-well culture plates (approx. 500,000 cells) or T25 tissue culture flasks (approx. 85% confluency) and cultured during experiments in sterile DMEM/F-12 with 1 mg/ml BSA at either 20% O_2_ or 2% O_2_. Cells were treated by adding to the culture medium 1-10 μg/ml α-amanitin (Sigma-Aldrich), 250 μM CoCl_2_ (Sigma-Aldrich), inhibitors (EDM Chemicals, Inc., Gibbstown, NJ) specific for MMP2 and MMP9 ((2R)-[(4-Biphenylylsulfonyl)amino]-N-hydroxy-3-phenylpropionamide, BiPS; 100 nM), MMP2 only ((2-((isopropoxy)-(1,1′-biphenyl-4-ylsulfonyl)-amino))-N-hydroxyacetamide; 250 nM) or MMP9 only (MMP9 Inhibitor I; 100 nM), specific inhibitors for ERK, JNK, and p38 (Jessmon et al., 2010), HSP70 inhibitor (VER-155008, Santa-Cruz Biotech; 0.05-100 μM), and 10 nM recombinant MMP2 (R&D Systems; rMMP2;). For protein analysis attached cells were extracted using cell lysis buffer (Cell Signaling) and total cellular protein concentrations were determined using Pierce BCA protein assay kit (ThermoFisher Scientific). The HTR-8/SVneo cell line was maintained in DMEM/F12 with 10% donor calf serum at 20% O_2_ between passages 30-50, and routinely checked for production of β-hCG, KRT7, and when cultured on Matrigel, HLA-G. Serum was replaced with BSA beginning 24 h before all experiments.

### Villous Explant Culture

Placental tissues were obtained with Wayne State University Institutional Review Board approval and patient informed consent from first trimester terminations at a Michigan family planning facility. Fresh tissue was placed on ice in PBS and immediately transported to the laboratory. The chorionic villi were dissected into pieces of approximately 5 mg wet weight and transferred individually into DMEM/F12 culture medium supplemented with 10% donor calf serum, 100 I.U. penicillin and 100 μg/ml streptomycin in a 24-well culture plate (Costar, Corning, NY) for villous explant culture (Bolnick et al., 2015). Treatments were performed as described for the cell line.

### Antibody Arrays

Cell lysates (1 mL) were incubated overnight with antibody array membranes, using a human MMP antibody array kit (Abcam). The membranes were washed and incubated with a secondary biotin-conjugated antibody, followed by incubation with horseradish peroxidase–conjugated streptavidin, according to the manufacturer’s direction. The arrays were developed, using enhanced chemiluminescence, and imaged on the ChemiDoc Imaging System (BioRad). Labeling of each protein in the array was quantified using ImageJ software (NIH – http://rsbweb.nih.gov/ij/). The mean of six negative controls was subtracted for background correction, and the mean of six positive controls was used to normalize the data and calculate the relative expression levels.

### Western Blotting

Western blots were performed as previously described (Kilburn et al., 2000). Briefly, cellular lysates were diluted in SDS sample buffer containing 5% β-mercaptoethanol, run on precast 4%–20% Tris-HCl gradient gels (BioRad), and blotted with antibodies against MMP2, MMP9, HIF1A (R&D Systems), HIF2A/EPAS1 (Novus Biologicals) and HSPA6 (Abcam) diluted 1:1000 in TTBS and 5% milk. Densitometry was used to quantify grey levels of protein bands of interest, using image analysis software (SimplePCI, Hamamatsu). Background grey levels, determined in a blank lane, were subtracted to obtain the specific grey level for each band.

### ELISA

Cells were cultured and treated in six-well plates. ELISA was conducted using HBEGF & MMP2 DuoSet ELISA Development kits (R&D Systems). The optical density of the final reaction product was determined at 450 nm using a programmable multiplate spectrophotometer (Power Wave Workstation; Bio-Tek Instruments) with automatic wavelength correction. Data are presented as nanograms (ng) of HBEGF or MMP2 per microgram (μg) of total protein, determined using standard curves generated with the respective recombinant proteins (R&D Systems).

### Immunocytochemistry

TB cells were labeled with monoclonal antibodies against HBEGF (R & D Systems) diluted 1:500, MMP2 (R & D Systems) diluted 1:200, HSPA1A (Abcam) diluted 1:400, or HSPA6 (Abcam) diluted 1:400. To visualize and quantify bound primary antibody, an Envision System peroxidase anti-mouse/rabbit kit (DAKO) was used. Staining (grey level) was imaged using a Leica DM IRB epifluorescence microscope, and images were captured using a Hamamatsu Orca digital camera. Images were semi-quantified using SimplePCI (Hamamatsu) imaging software, as previously described (Leach et al., 2012).

### qPCR

RNA from HTR-8/SVneo cells was collected using the miRNeasy kit (Qiagen), according to the manufacturer’s protocol. RNA concentration was determined using the NanoDrop spectrophotometer and purity was ascertained with a microfluidic Bioanalyzer (Agilent Technologies - 2100 Electrophoresis Bioanalyzer Instrument). RNA was used in subsequent qPCR (Bustin et al., 2009) analysis. Reverse transcription was performed using the Quantitect Reverse Transcription kit (Qiagen), and qPCR for MMP2, HBEGF and HSPA6 was conducted with the Quantitect SYBR Green PCR kit without UNG (Qiagen), in a final volume of 25 μl. GAPDH was used as a housekeeping gene to normalize the data. Semi-quantiative analysis was performed according to the ΔΔCt method (Pfaffl, 2001). Primers for GAPDH, HBEGF and MMP2 were obtained from Qiagen.

### LongRNA (LnRNA) Library Prep for Next Generation Sequencing (NGS)

LnRNA was isolated using the miRNeasy mini kit (Qiagen). The RNA was quantified and its purity assessed with an Agilent 2100 Electrophoresis Microfluidics Analyzer. RNA was converted into an adapter-ligated cDNA library, using Ovation & Encore Library Preparation Kit (NuGen), according to the manufacture’s protocol. Each cDNA sample was bar-coded, and the resulting 12 libraries (three replicate libraries for each of the four O_2_ conditions) were combined for NGS. Paired-end sequencing was performed for 50 cycles using the Illumina HiSeq-2500 sequencer.

### Data Alignment & Mapping

RNA sequencing data was first processed with demultiplexing software (Casava 1.8.2, Illumina). It was then aligned to the human genome build HG19, and to the ribosomal sequences 18s and 28s, using bioinformatics tool Novoalign (Novocraft, 2010). Novoalign determined unique alignments that were used to generate 1000 reads per coding segment per sample. The reads thus generated were converted into bed.files and imported to the Genomatix mapping station (GMS) (**Genomatix Software GmbH**). The GMS, using RNA-seq analysis, generated data in the form of Reads Per Kilobase of exon per Million fragments mapped (RPKM) for 25,000 genes in the database.

### Promoter Extraction

The Genomatix Genome Analyzer (GGA) MatInspector program (Cartharius et al., 2005) was used to extract the promoter regions for the differentially regulated genes.

### Statistics

All statistics were performed with GraphPad Prism 6 software. One-way ANOVA with Student–Newman–Keuls *post hoc* comparisons was used to identify changes between controls and treatments. Two-way ANOVA was used for experiments with multiple groups. Significance was defined as *P* < 0.05; all data are expressed as mean ±SD.

## ACKNOWLEDGEMENTS

We thank Northland Family Planning Centers of Michigan and their patients for participating in this research study. The authors are grateful to Anelia Petkova for technical assistance, and to Dr. Stephen A. Krawetz, Dr. Susan Dombrowski, Robert Goodrich, and Edward Sendler for training in next-generation sequencing library preparation and bioinformatics.

## AUTHORS CONTRIBUTIONS

The study and experimental design was conceived by DRA. CVJ, PJ, CTB, ADB and BAK performed the experiments and contributed to data analysis. CVJ and DRA wrote the manuscript. MH provided placental tissue. All authors discussed the results, and contributed modifications of the manuscript.

